# iNaturalist mammal observations classified by evidence type

**DOI:** 10.1101/2025.09.30.678090

**Authors:** Mohammad Alyetama, Alex J. Jensen, Yakira Jackson, Jane Widness, Roland Kays, Benjamin R. Goldstein

## Abstract

Ecologists show growing interest in observational data generated by participatory scientists. For mammals, the largest participatory science platform is iNaturalist, which has more than 2.8 million observations from North American represented through images of living animals, dead animals, tracks, and scat. These different types of evidence could give insight into the underlying sampling paradigm for an observation (e.g. dead animals might be more likely to be reported near roads) and thus may be useful for scientific applications of these data. However, while iNaturalist allows users to annotate observations by evidence type, many observations are not annotated. We use machine learning (ML) to classify the evidence types associated with observations of North American mammals in iNaturalist, adding metadata that can be used to subset data or to model multiple observation processes. Here, we present a dataset containing metadata augmenting 1.33 million North American mammal iNaturalist observations with evidence type. Each observation is categorized as either live animal, dead animal, tracks, scat, or other sign, and an associated confidence score is provided.

## Introduction

Ecologists are increasingly interested in analyzing unstructured participatory science data, both as a primary dataset documenting animal occurrences [1] and as a supplemental dataset in integrated models [2]. For mammals, the largest such dataset is iNaturalist, which has accumulated nearly 4 million “research grade” observations of mammals as of January 2025 (iNaturalist.org). iNaturalist sampling is unstructured, with no reporting of sampling process, spatial and temporal allocation of effort [3,4], the taxonomic preferences of individual users [5– 7], and logistical factors limiting detectability and species identification [8,9]. Robust inference on biological processes using these data requires accounting for these sources of sampling heterogeneity, which could otherwise bias results [4]. Consequently, the use of these data is limited by ecologists’ capacity to detect sampling heterogeneity and subsequently employ appropriate data filters and statistical models.

One important source of observational variation is evidence type. iNaturalist research grade observations are required to include a piece of evidence—typically a photograph or audio recording—that can be used by other members of the online community to verify the species ID [10]. While many iNaturalist observations of mammals have photos of live animals, many others are photos of dead animals such as roadkill; photos of animal tracks and scat; or photos of markings, nests, or burrows. These evidence types may follow different spatial patterns in their effective effort, meaning that species more frequently associated with one evidence type will be detected at a higher rate where effort for that type is higher. For example, dead animals may be more frequently detected in areas with more roads, while elusive but charismatic mammals with distinctive footprints such as gray wolves (*Canis lupus*) may be reported at a higher rate in regions with substrate that holds animal tracks (e.g., snow, mud). Models that assess the distribution of reports of those species without considering evidence type could wrongly conclude that populations are at higher densities in regions conducive to track-based identification. In addition to improving statistical inference, categorizing evidence types could also help in identifying human observers of certain archetypes, such as trackers or casual observers. The iNaturalist platform allows users to add evidence type annotations to observations, such as “dead” or “tracks,” and the iNaturalist community increasingly adds such annotations to existing records, but many observations remain un-annotated.

Machine learning-based image classification algorithms offer a solution to this problem. Machine learning-based image classification has become a powerful tool in ecological research, enabling automated species identification and behavioral annotation from photographs at large scales [11]. Convolutional neural networks (CNNs), in particular, have demonstrated strong performance in classifying wildlife images from camera traps [12]. These models learn visual features directly from training data and can generalize to complex, variable conditions such as lighting, angle, or background clutter. In participatory science contexts like iNaturalist, ML-based classification offers a scalable solution to annotate large datasets with metadata such as evidence type.

Here, we provide evidence type metadata for 1,326,100 observations of North American mammals submitted to iNaturalist. We trained a YOLOv8x image classification model [13] using manually labeled data to predict one of five evidence types: live animal, dead animal, tracks, scat, or other sign (Figure 1). We provide both the dataset and a reproducible pre-trained model that can be easily applied to classify new data. These outputs represent the first and most critical step toward downstream analyses of iNaturalist data that address heterogeneity in evidence types.

**Figure 1.**
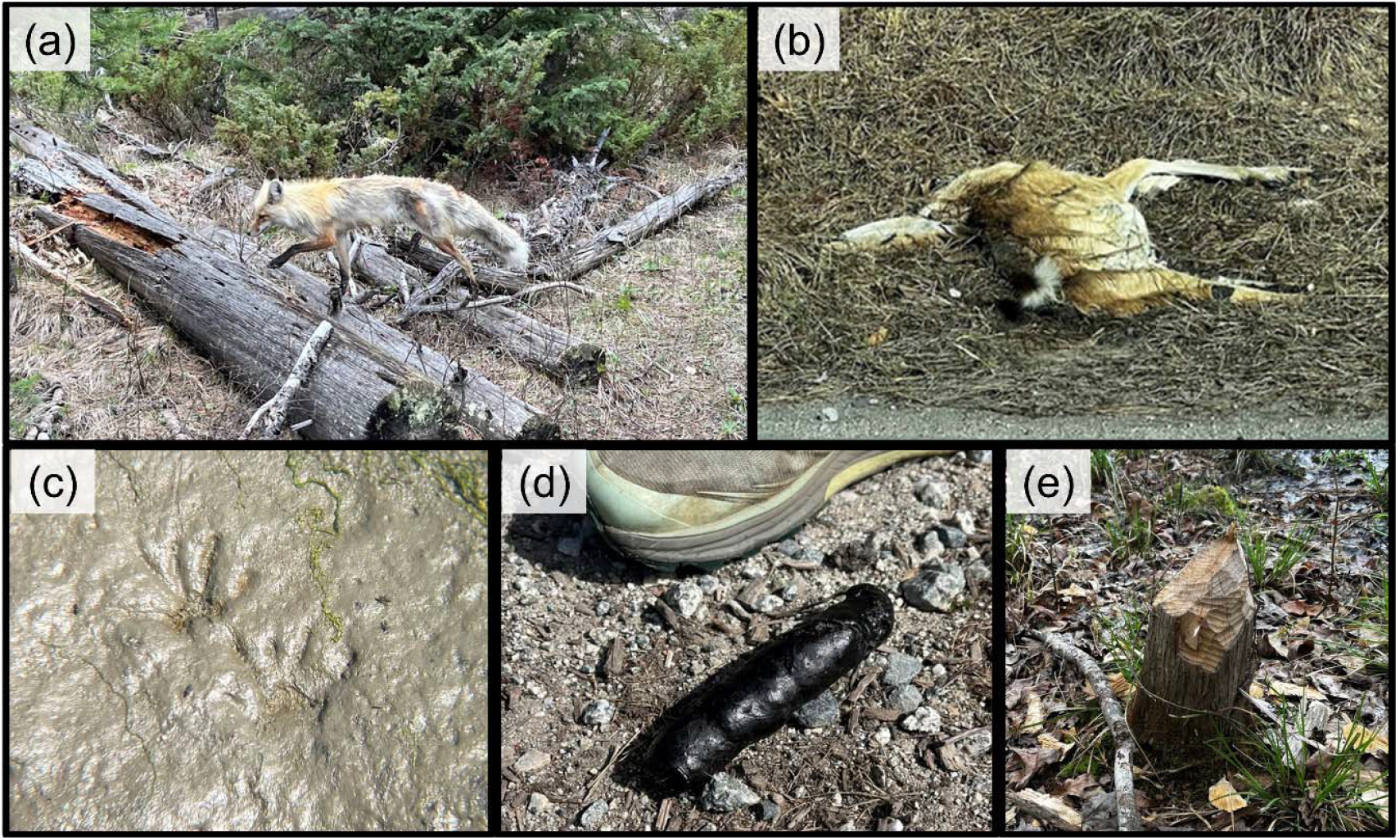
Example photos representing the five primary iNaturalist evidence type classes: (A) a live animal; (B) a dead animal; (C) an animal track; (D) animal scat; and (E) other sign, in this case tooth marks left on a tree stump. All images are obtained from iNaturalist under a CC0 license; photos A, C and D are attributed to Benjamin R. Goldstein and photos B and E are attributed to Michael V. Cove.

## Results

We provide evidence type classifications for 1,326,100 iNaturalist observations of North American mammals. The data are provided as a single large .csv file containing one row for each observation. For each observation, we provide the internal iNaturalist observation ID; the URL at which the image is hosted; the English common name and scientific name of the species; the date and time of the observation; the latitude, longitude, and positional accuracy of the observation; and a Boolean indicating whether the position of the observation was obscured. We maintain the column names to match the format of an iNaturalist data download. The dataset is published on Zenodo under the DOI 10.5281/zenodo.17177962. The trained classification model is publicly available on Hugging Face at huggingface.co/spaces/Alyetama/AiNaType, where users can interact with a live demo to test the model on their own images. The model weights, along with the full training and inference code, are hosted on GitHub at https://github.com/Alyetama/AiNaType. Detailed instructions for applying the model to other datasets, including custom iNaturalist exports, are provided in the repository’s README file.

Across all observations, 97.4% were associated with a confidence score in the top class of at least 0.8, 95.9% had a top confidence score of at least 0.9, and 91.2% had a top confidence score of at least 0.99. We classified 85.0% of all observations as live animals, 7.3% as dead animals, 1.6% as scat, 4.1% as tracks, and 1.9% as other sign (Figure 2B). Classifications varied widely by taxonomic order, with observations of primates comprising 99.3% live animal photos but observations of Eulipotyphla (shrews and moles) comprising 84.9% dead animals (Figure 2C).

**Figure 2.**
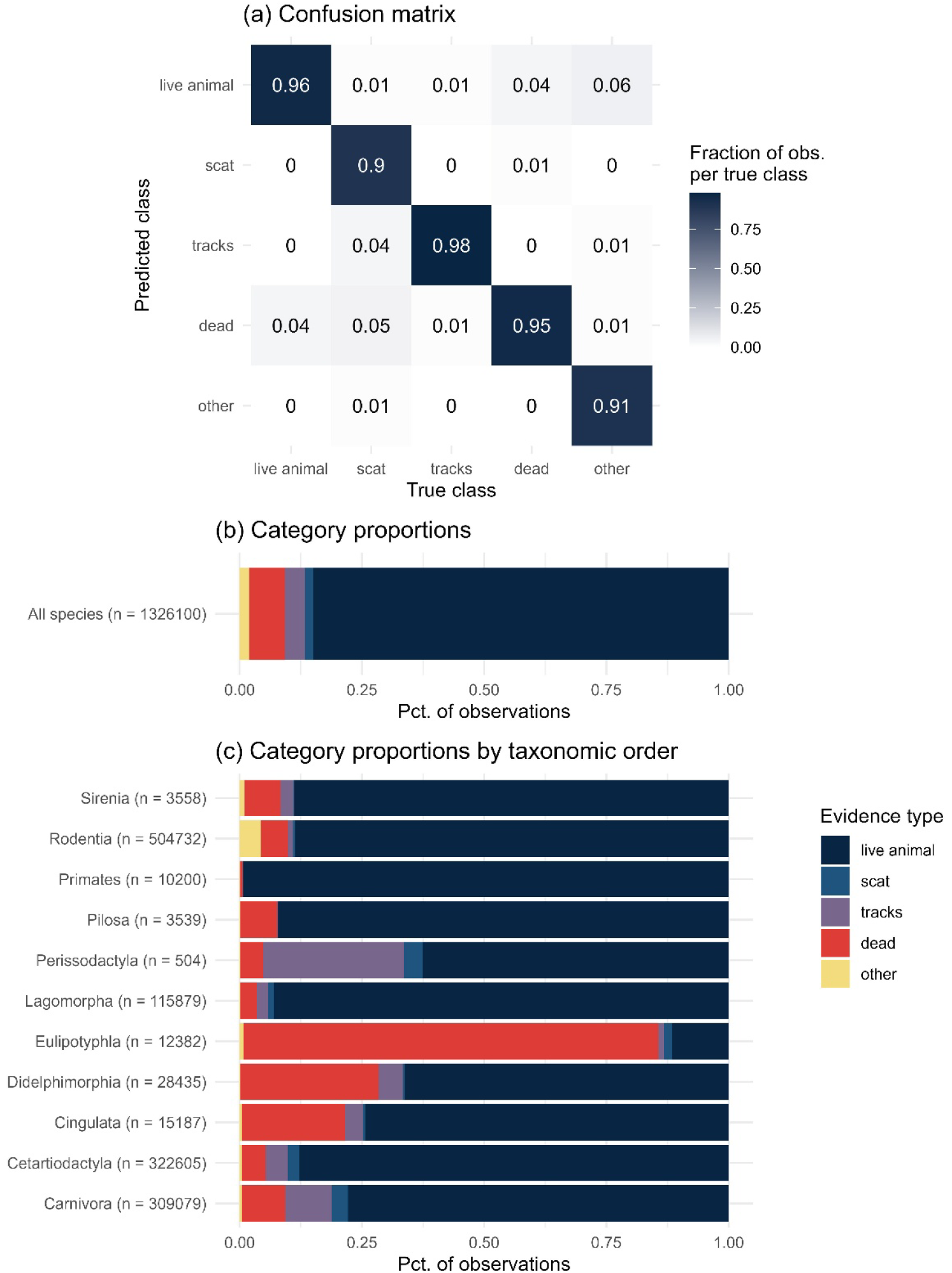
(a) Normalized confusion matrix for the AiNaType-v8x model on the validation set. The model shows high accuracy across all evidence types. (b) Overall rates of five evidence types. (c) Rates of each evidence type by taxonomic order.

We cross referenced a random subsample of our classified data against existing iNaturalist annotations. Among 2,000 observations we classified as each of dead animals, tracks, or scat, 44.6%, 34.3% and 28.1, respectively, were annotated as such in iNaturalist. Moreover, only 12.3% of observations of living animals were annotated as such.

## Discussion

Our dataset represents the first attempt to classify iNaturalist observations by evidence type in a scalable way. This procedure has yielded useful metadata that can enhance the downstream analysis of more than 1 million participatory science records. These data can be used to filter iNaturalist data to meet more stringent data standards—for example, in an analysis highly sensitive to false positive errors in the data, it may be advantageous to exclude scat detections of a species, which may be prone to misclassification [14]. These data could be used as covariates in distribution models to represent how a species may be more or less detectable depending on the sampling method. Further study is needed to understand the implications of various representations of evidence type in a statistical model. Finally, these data support analyses of individual, regional, temporal, or cultural differences in how participatory scientists engage with the natural world, shedding light on the impact of iNaturalist on the human-wildlife relationship [5]. These data could also serve as a reference for community members who want to quickly find sign observations within iNaturalist to add annotations after manually confirming the classification.

We provide confidence scores for each classification (0-1, summing to 1 across categories) and recommend that practitioners consider filtering observations by confidence level for their applications, such as only including observations with a confidence score above 0.9. The appropriate threshold will depend on the application, balancing sensitivity to false positives (for which there will be fewer if a higher threshold is used) with data requirements (since lower thresholds will admit more total observations).

The model has some difficulty distinguishing between truly dead animals and those that only appear dead due to visual context. For example, opossums may mimic death as a natural defense behavior, leading to misclassification. Similarly, bats, small rodents, and shrews are often photographed while being held in human hands, where their limp posture can resemble that of a dead animal. These scenarios can confuse the model, which relies solely on visual cues without access to behavioral or contextual information. Another potential discrepancy is our choice to classify animal parts, such as shed antlers, in the “dead” category along with bones, although the animal may still be alive. Additionally, because we classified only the first observation of each image, there may be additional classification errors for cases where additional photos contain other evidence, such as an iNaturalist observation that contains both a photo of tracks and a photo of scat.

## Materials and Methods

### iNaturalist data curation

For training, we batch-exported data from iNaturalist.org in January 2023 by filtering for observations of mammals in North America from January 1, 1900 to December 31, 2022. This initial dataset contained 1,512,795 unique research grade observations. We used each record’s observation ID to download metadata from the iNaturalist Open Dataset (iNOD) via the Registry of Open Data on Amazon Web Services [15]. The iNOD only includes observations with images licensed under Creative Commons or Public Domain terms (CC0, CC-BY, CC-BY-SA, CC-BY-NC, CC-BY-NC-SA, CC-BY-ND, and CC-BY-NC-ND). We then filtered our observation dataset to retain only observations for which the primary photo was available through iNOD. We downloaded the photos using a custom Python script designed to retrieve and organize the photos. After manually developing the training dataset, we updated our full dataset by retrieving all records through February 27, 2024, following the same method described above.

We accessed a targeted subset of annotated data to quickly amass training images representing the four less common evidence types (dead animal, tracks, scat, other sign) that we then manually reviewed. We used the iNaturalist API (https://api.inaturalist.org/v1/docs/) to access annotations on observations that were not available within the iNaturalist export tool or the Global Biodiversity Information Facility at the time (note that the iNaturalist GBIF product now includes annotations). For the dead animal category, we separately downloaded observations with the annotations ‘Alive or Dead: Dead’ and ‘Evidence of Presence: Bone’. For tracks and scat we used ‘Evidence of Presence: Track’ and ‘Evidence of Presence: Scat’, respectively. We also exported all of the observations from the North American Animal Tracks Database, which is a project within iNaturalist that contains observations with both tracks and sign evidence types. For the other sign category, we subset our observation data to include species that commonly make other types of sign: pocket gopher (*Thomomys spp*., mounds of dirt); beaver (*Castor canadensis*, chew marks on trees, dams); black bear (*Ursus americanus*, scratching); woodrat (*Neotoma spp*., stick nests); armadillo (*Dasypus novemcinctus*, burrow); gray squirrel (*Sciurus carolinensis*, nests); deer (*Odocoileus spp*., tree rubs); and badger (*Taxidea taxus*, burrow).

After classifying the full dataset, in July 2025 we separately queried the iNaturalist API for annotations associated with 2,000 random observations that we classified as each of live animals, dead animals, tracks, and scat. We recorded the rate at which annotations in iNaturalist matched our classifications and recorded the fraction of observations that had any annotations at all.

### Development of training and validation data

We used Label Studio, a self-hosted open-source annotation platform, to manually annotate images for training and validation. Each image was independently reviewed and assigned to one of the five evidence type classes. In total, we manually annotated 64,120 images, including 22,301 labeled as live animals, 24,105 as dead animals, 9,193 as tracks, 4,077 as scat, and 4,344 as other. When necessary, annotators cross-checked associated metadata and additional images from the same observation to improve labeling accuracy. To minimize bias, multiple annotators were involved in labeling, and disagreements were resolved through discussion or consensus review. We then split the labeled dataset into training and validation sets, stratifying by evidence type to preserve the distribution of classes across both subsets. We used 80% of the labeled data for training and reserved 20% for validation to assess model performance.

During the data curation process, we excluded some images (1,179) from the training dataset due to issues such as heavy blurring, severe underexposure, or photos of computer screens displaying images rather than direct wildlife observations, which were unlikely to provide useful visual features for training. Additionally, during early iterations of training, we observed a substantial number of bat (Chiroptera) observations being classified as “dead” even when the animal was alive. This was because a large proportion of bat images showed individuals being held in human hands, often appearing lifeless due to the posture or context. To avoid introducing systematic confusion and mislabeling, we opted to exclude all bat species from the final dataset used for model training and inference.

### Machine learning classification

We trained an image classification model to categorize iNaturalist mammal observations by evidence type using the Ultralytics YOLOv8 framework (version 8.2.48), specifically the YOLOv8x-cls architecture with 133 layers and approximately 56 million parameters. Training was conducted on an NVIDIA GeForce RTX 4090 GPU using PyTorch 2.3.1 with CUDA acceleration, using a dataset of 64,120 labeled images split into stratified training and validation subsets. Key training parameters included an image size of 224 × 224 pixels, batch size of 128, up to 100 epochs with early stopping after 50 epochs without improvement (best model achieved at epoch 79), an initial learning rate of 0.01 with a learning rate factor of 0.01, momentum of 0.937, weight decay of 0.0005, and stochastic gradient descent (SGD) as the optimizer. We applied data augmentation strategies, including horizontal flips (0.5 probability), HSV adjustments (hue: 0.015, saturation: 0.7, value: 0.4), random translations (0.1), and erasing (0.4), as well as Mosaic and RandAugment policies to improve generalization. Mixed precision training (AMP) was enabled to accelerate training, and deterministic mode was set to ensure reproducibility.

### Performance metrics

Model performance was evaluated on a held-out validation set using standard classification metrics. The final YOLOv8x-cls model, which we named AiNaType-v8x, achieved a top-1 classification accuracy (the percent of observations for which the top predicted category matched the true category) of 95.01%. A confusion matrix was generated to assess the model’s ability to distinguish among the five evidence types (Figure 2A), which confirmed strong class-wise performance. The model struggled most at differentiating between live and dead animals, but still only misclassified dead animals as live and vice versa 4% of the time.

## Acknowledgements

Thanks to Michael V. Cove for providing example photos used in Figure 1. Thanks to the North Carolina Museum of Natural Sciences Biodiversity Lab for resources and support. Funding was provided by NSF Awards 2206783 and 2206784.

